# Fruit colour and range size interact to influence diversification

**DOI:** 10.1101/2021.10.26.465838

**Authors:** Adrian P. Hill, Maria Fernanda Torres Jiménez, Nicolas Chazot, Cibele Cássia-Silva, Søren Faurby, Christine D. Bacon

## Abstract

**Aim:** Different fruit colours are associated with dispersal by different frugivores, largely based on colour vision type. Frugivore mobility affects overall range size for the plant being dispersed. Here we determine the interaction between different fruit colours, range sizes, and diversification rates by testing two hypotheses: That (1) fruit colours attractive to birds have larger range sizes due to their higher dispersal ability, and that (2) different frugivore disperser groups, bird or mammal, leads to different diversification rate at different range size, where intermediate range size leads to the highest diversification rate.

**Location:** Global.

**Time period:** Contemporary (or present)

**Major taxa studied:** Palms (Arecaceae)

**Methods:** Using model selection, we identified three groups of colours with similar diversification rate and likely disperser. Range sizes were estimated and categorized species as small, intermediate, or large-ranged. For model selection and to determine the relationship beween fruit color, range size and diversification rate we used Multi-State Speciation and Extinction (MuSSE) models.

**Results:** Species with intermediate range size had the highest net diversification for all three fruit colour groups. Bird-dispersed palms more likely diversified at small than at large range size while mammal-dispersed palms more likely diversified at larger range size than small. Fruit colours associated with mammal dispersal had more large-ranged species than colours associated with bird dispersal.

**Main conclusions:** The associated between intermediate range size and higher diversification rate indicates that spatial factors that affect diversification at small and large range sizes result in higher diversification at intermediate ranges. We find striking differences in diversification rate within each range size category between fruit color groups. This suggests that the relationship between diversification rate and range size depends on the specific frugivorous dispersers and their dispersal patterns. This study reveals how fruit traits alter dispersal patterns and how that, in turn, influences diversification.

## 1 INTRODUCTION

In order to disperse, plants produce fruits for seed ingestion by frugivorous animals. To increase the chance of consumption, and thereby dispersal, fruits exhibit colours that increase the probability of detection by desirable frugivores and reduce detection by nondesirable ones (Melo, Penatti, & Raizer, 2011). Animals better detect fruits that contrast against background colours, for example red fruits against green foliage for tri- and tetrachromatic frugivores. Bird and mammal frugivores are the most significant seed dispersers in terrestrial habitats (Fleming & Kress, 2011), but due to variation in colour vision these groups perceive fruit colour contrasts differently.

Birds are tetrachromats and have the ability to discriminate between red and green (Vorobyev et al., 1998), whereas mammals are mostly dichromats and do not distinguish between red and green (Jacobs, 1993), except for some primate groups (apes, old world monkeys, and a few new world monkeys; Onstein, et al., 2020; Regan, et al., 2001). Birds are therefore more likely to consume fruits that are red, black and intermediate shades of purple (Duan, Goodale, & Quan, 2014; Schaefer, Valido, & Jordano, 2014). These fruits are often collectively termed bright in the literature, and may include orange and yellow (Onstein et al., 2019; Knight & Siegfried, 1983). Fruit colours equally detectable and often preferred by mammals are often termed dull and include green, brown, orange, and yellow fruits (Janson, 1983; Sinnott-Armstrong et al., 2018). Whether it is birds or mammals that more likely consumes any specific orange or yellow-coloured fruit may be determined by the brightness of the colour or whether it resembles more clearly delineated bird- or mammal-dispersed fruit colours. For example, orange, often being similar in appearance to either red, yellow or brown, could be consumed by frugivores depending on which of these colours it resembles more.

Frugivory-related traits such as fruit colour are important drivers of angiosperm diversification (Lu et al., 2019; Onstein et al., 2018, 2020). However, the mechanisms by which particular fruit colours influence diversification remain unclear. One potential mechanism is that certain fruit colours promote diversification due to their synergic effects with geographic range (Lu et al., 2019; Onstein et al., 2019). The specific dispersers associated with a plant will determine its dispersal ability, thus, the fruit colour used to attract one disperser or the other can be an important predictor of plant geographic range. Overall, birds travel and disperse seeds further (Santos, Tellería, & Virgós, 1999; Stevenson, Cardona, Cárdenas, & Link, 2021), and cross geographical barriers with greater ease than (non-flying) mammals do (Lu et al., 2019). Therefore, plants that produce red, black, and purple fruits often disperse further due to their association with bird dispersal (Lu et al., 2019), resulting in a wider geographic range for that plant species.

Different mechanisms that facilitate or hinder speciation and extinction are associated with different range sizes (Gaston, 1998). Furthermore, range size is correlated with dispersal ability (Estrada et al., 2015; Faurby & Antonelli, 2018; Penner & Rödel, 2019; Sinnott-Armstrong et al., 2018). Therefore, plants dispersed by animals with high dispersal ability may have larger range sizes, and be influenced by mechanisms impacting speciation and extinction at large range size (Bacon et al., 2013). Geographically isolated populations are more likely to speciate and are typical of species with high dispersal ability as these may colonise new areas (Lester, Ruttenberg, Gaines, & Kinlan, 2007). For example, a large range size may facilitate parapatric speciation through an isolation by distance effect (Baptestini, De Aguiar, & Bar-Yam, 2013), or ecological speciation due to variation in environmental conditions (Bacon et al., 2013; Chen & Schemske, 2015; Keller & Seehausen, 2012) or differences in pollinator communities (Neves et al., 2020). However, high dispersal ability may also hinder diversification through the maintenance of gene flow within a species range (Claramunt, Derryberry, Remsen, & Brumfield, 2012).

Diversification rate may, alternatively, be higher in small- or intermediate-ranged species, which have geographic barriers embedded within the range (Gaston, 1998). Thus, connectivity around the barrier for species unable to cross would hinder allopatric speciation. However, small-ranged species are more prone to be driven to extinction, which impacts net diversification (O’Grady, Reed, Brook, & Frankham, 2004). Small-ranged species may not benefit as often from the speciation-facilitating factors suggested for large-ranged species such as ecological speciation. Owing to the hindrances of diversification associated with either range size extremity, intermediate ranges are expected to balance gene flow and isolation, and thus lead to the highest diversification rate.

With a wide variety of fruit colours, size, shape, and amount of fruits, palms are a keystone resource for frugivorous animals (Zona & Henderson, 1989). Palms rely on a wide array of animals (e.g. birds, bats, non□flying mammals, reptiles, insects, and fishes) to disperse their seeds (Zona & Henderson, 1989). This mutualistic interaction has been important in shaping palm distribution patterns over space and time (Lim, Svenning, Göldel, Faurby, & Kissling, 2020; Onstein et al., 2018; Sales, Kissling, Galetti, Naimi, & Pires, 2021). Here, we test the impact of the interplay between fruit colour on range size and diversification rate in palms (Arecaceae). Palms are a taxonomically (c. 2,600 species; Baker & Dransfield, 2016) and functionally diverse clade (Kissling et al., 2019) characteristic of tropical regions (Couvreur, Forest, & Baker, 2011). Palms have rich phylogenetic (Faurby et al., 2016), trait (Kissling et al., 2019), and distribution data (GBIF; https://www.gbif.org/) available. Taken together, palms are an excellent case to study the potential interaction between fruit colour and range size on diversification rate.

Here, we hypothesise that (1) large range size is associated with fruit colours that are dispersed by frugivorous birds, owing to their mobility and high dispersal ability. We also hypothesise (2) that net diversification varies between fruit colour groups associated with different dispersers, revealing how dispersal ability impacts diversification at different range sizes. Due to the potential effects that hinder speciation at either range size extreme (small and large), we expect that diversification rate is higher for palms with intermediate range size.

Plants have dynamic dispersal systems in that a single individual may rely on a multitude of different animal species to disperse its seeds. These interactions have shaped global plant diversity and biogeography. It is therefore important to examine how the colour of a fruit ultimately leads to more diverse lineages through through time and has large-scale impacts on biogeography and evolution.

## 2 MATERIALS AND METHODS

### 2.1 Data

We obtained fruit colour data for 1,485 palm species (ca. 57% of all recognized species; Baker & Dransfield, 2016) from the PalmTraits 1.0 database (Kissling et al., 2019). For phylogenetic analyses, we used a set of 30 trees sampled from a posterior distribution (an updated version of the phylogeny from Faurby et al., (2016) which is described in Data Availability). Species occurrence records for 1,785 species were obtained from the Global Biodiversity Information Facility (last consulted on 31 January 2019; GBIF Occurrence Download https://doi.org/10.15468/dl.rjmqfy). We performed all analyses using R, version 3.6.3 (R Core Team, 2019).

### 2.2 Model selection

To estimate how different fruit colours affect speciation and extinction rate of palms, we used Multi-State Speciation and Extinction (MuSSE) models (‘diversitree’, version 0.9-13; FitzJohn, 2012). We removed all polymorphic species from the data (i.e. species with more than one fruit colour assigned). Then, we classified the remaining species into five fruit colour categories based on likely frugivorous disperser group (bird or mammal): 1) black, purple; 2) red, orange; 3) yellow; 4) brown, green, blue; 5) white. Species with ambiguous colour definitions from PalmTraits: “ivory”, “straw-coloured”, “cream”, “pink”, and “grey” (70 species total) were excluded.

Instead of only fitting MuSSE models with the five categories, we tested which categories could be merged together without significant loss of model fit. Starting with the most complex model (five colour categories), we tested all ten possible combinations of merged categories from the five initial categories. In each case two of the initial five groups were combined while keeping the other three seperate. All models were fit on 30 trees sampled from the posterior distribution of trees. To compare AIC scores between models, we calculated a median ΔAIC score across trees by subtracting the AIC scores for each model (values from each of the 30 models on 30 trees) by the AIC scores of the full five-category model and then calculating the median of these ΔAIC values. The best model (highest ΔAIC) from the first round (which had four categories per model) was selected and used as a starting point for the second round of merging. The best model from this round (which had three categories) was tested in a third round to check if additional simplification was possible, but these produced ΔAIC scores that dramatically reduced model fit.

The best model had the following fruit colour categories: 1) black, purple, red, orange, white; 2) yellow; and 3) brown, blue, green. Based on the likely most frequent frugivorous dispersers for the selected model fruit colour groups we hereafter refer to the groups as 1) bird, 2) mammal-1, and 3) mammal-2.

With the selected model, there were some species from the original dataset which, based on the colour combinations in the selected model were no longer polymorphic as their colours were now grouped into a single category. We added these species back into the fruit colour dataset for the remaining analyses. This resulted in 1063 palm species with fruit colour data, representing *ca.* 41% of all recognized palms species (Baker & Dransfield, 2016).

### 2.3 Range size calculations

We estimated the range size of all species with available, filtered data. Fossil data, records without coordinates, duplicates, and occurrence records nearby science institutions, within bodies of water or country centroids with a buffer of 5000m were filtered out using ‘CoordinateCleaner’ 2.0-3 (Zizka et al., 2019). We also removed records outside the species native “botanical country”, which is the level three geographic division defined by the International Working Group on Taxonomic Databases for Plant Sciences (TDWG) (Brummitt, Pando, Hollis, & Brummitt, 2001), using a custom Python script (https://github.com/mftorres/palm_leaf/blob/master/notebooks/GBIF_variables_data_prep.ipynb).

We built alpha hull polygons to estimate the range (extent of occurrence) for species with three or more records using ‘rangeBuilder’ version 1.5 (Rabosky et al., 2016), keeping only terrestrial range (clipping to land). The package optimises the alpha value to create the smallest possible polygon(s) for each species. Because alpha hull polygons allow for concavities in the outer perimeter of the species range they are a more conservative range estimate compared to convex hulls (Meyer et al., 2017). Alpha hull area was calculated in ‘raster’ version 3.0-12 (Hijmans, 2020). For our analyses we did not need precise areas for all species but only the relative sizes (grouped into small, medium, and large range). We were therefore able to include relative range size for some species with fewer than three records if available data suggested that they have small ranges. More specifically we included species that occur exclusively within a single TDWG unit and with fewer than three occurrence records in that unit. These species were added into the range size dataset with an arbitrary range size that was lower than any of the alpha hull areas (0.1 km^2^), meaning they were categorised as small-ranged.

Species were then ordered based on ascending range size and then split into three range size categories: small, intermediate, and large. Each range size category contained a third of the species each. Species in these range size categories were further split by their associated fruit colour group into nine total groups based on both fruit colour and range size.

### 2.4 Relationship between fruit colour and range size on diversification rate

We built MuSSE models based on the nine combined categories of range size and fruit color (fruit colour and range size categories are mapped on a palm phylogeny in Figure 1). In order to simplify the model and reduce overfitting we disallowed changes between categories that would that would effectively mean two character transitions at once. This means transitions were allowed only within the same range size category for transitions that occurred between different fruit colour groups, or within the same colour group for transition between range sizes. Transitions between range sizes could only occur between adjacent ranges at one time, small-medium (and vice versa), medium-large (and vice versa). All other parameters were unconstrained.

**Figure 1.**
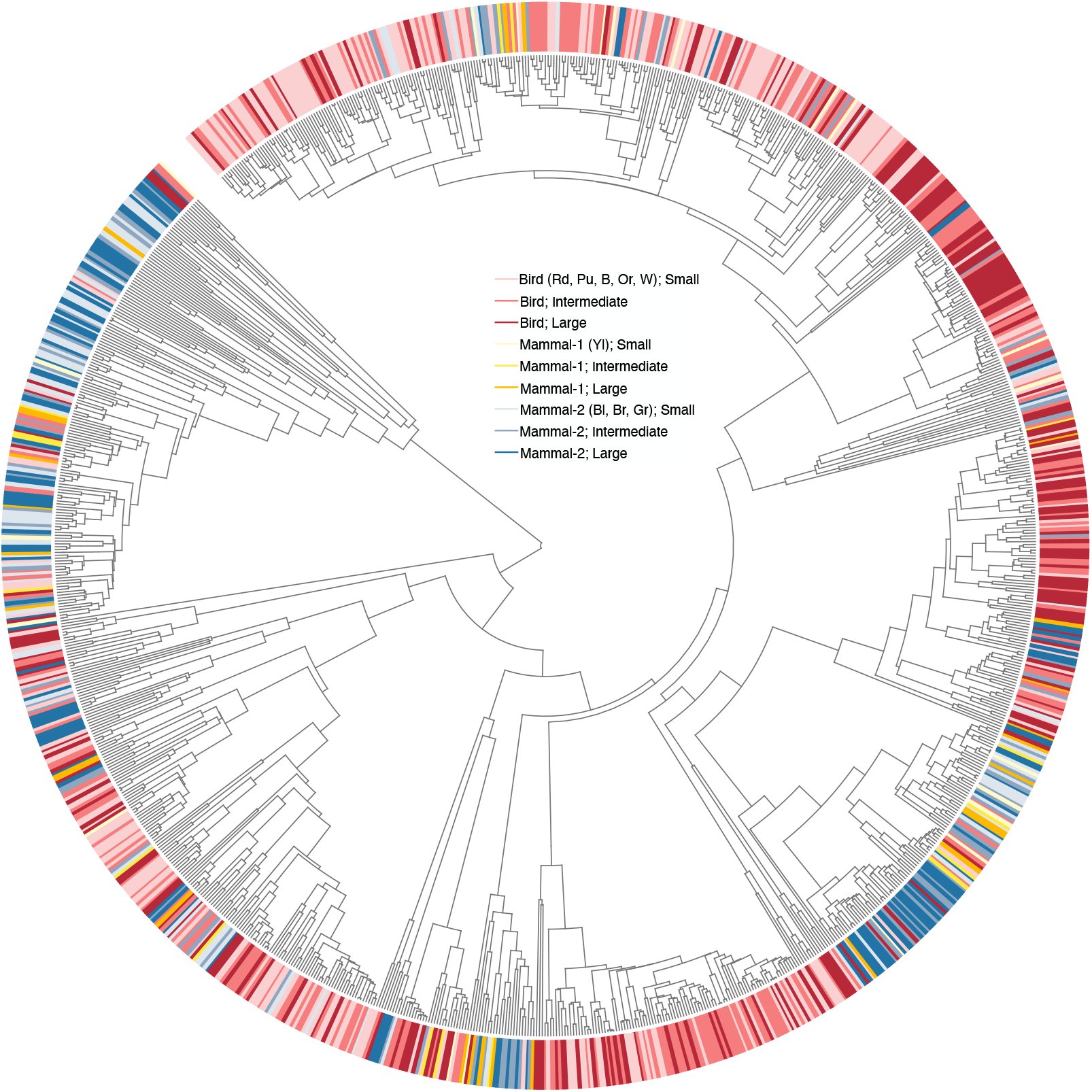
Fruit colour categories mapped onto the palm phylogenetic tree. Colour categories description. Bird group: Rd = red, Pu = Purple, B = black, Or = orange, W = white; Mammal-1 group: Yl = yellow; Mammal-2 group: Bl = blue, Br = brown, Gr = green.

## 3 RESULTS

For both the mammal-1 and mammal-2 palm groups, we found that large-ranged species make up the largest proportion of species (representing 42.4% and 39.9% of the respective groups; Figure 2). The mammal-1 group had more small- than intermediate-ranged species (33.3% small versus 24.2% intermediate), whereas the mammal-2 group had more intermediate- than small-ranged species (28.7% small versus 31.4% intermediate). For bird-dispersed palms, we found a roughly equal proportion of small and intermediate-ranged species (35.4% small versus 34.9% intermediate), whereas 29.7% of species were large-ranged. Based on a Fisher’s exact test (Mehta & Patel, 1983) we found that disperser group and range size was not independent (p=0.0089).

**Figure 2.**
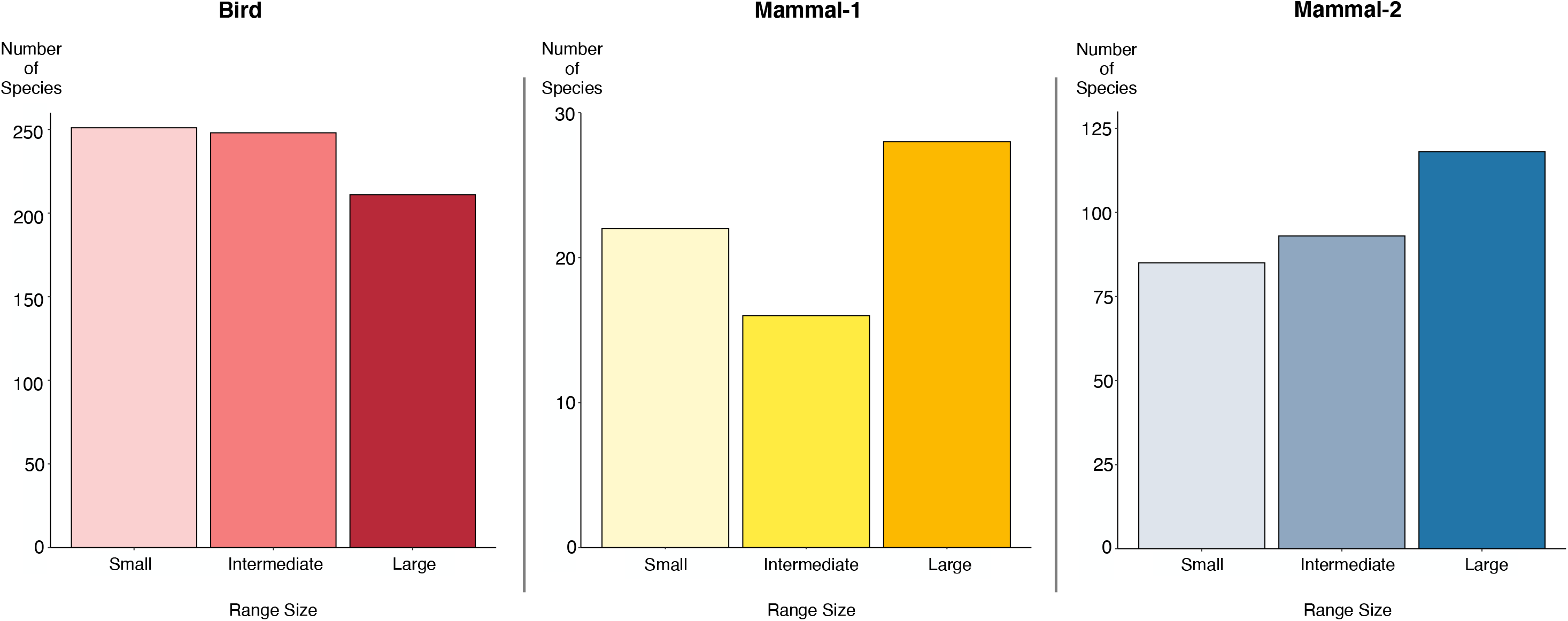
Number of species classified as small, medium, or large range within each fruit colour group.

The highest diversification rate was associated with the mammal-1-dispersed group, followed by the bird-dispersed and lastly the mammal-2-dispersed groups (Figure 3). We also identified a combined effect of range size and fruit colours on palm diversification rate. The net diversification rate was highest at intermediate range size (median ± 16.5% of species ranges) for all three fruit colour categories (Figure 3). Net diversification rate was higher for large-ranged than small-ranged bird-dispersed species and barely so for the mammal-1 fruit colour group. If the shape of the relationship were symmetrical, as the distribution shape within each fruit colour group may suggest (Figure 3), a higher net diversification for smaller-ranged than large-ranged species would indicate that the point of maximised diversification is lower than the median intermediate-ranged species (see Appendix S1 in Supporting Information). Small range size was associated with higher diversification than the large range for bird-dispersed fruit colour group (Figure 3), so for a symmetrical relationship this would indicate that the point of maximised diversification is smaller than exact intermediate for bird-dispersed species. For mammal-dispersed species the opposite is true, the point of maximised diversification is larger than exact intermediate (Figure 3).

**Figure 3.**
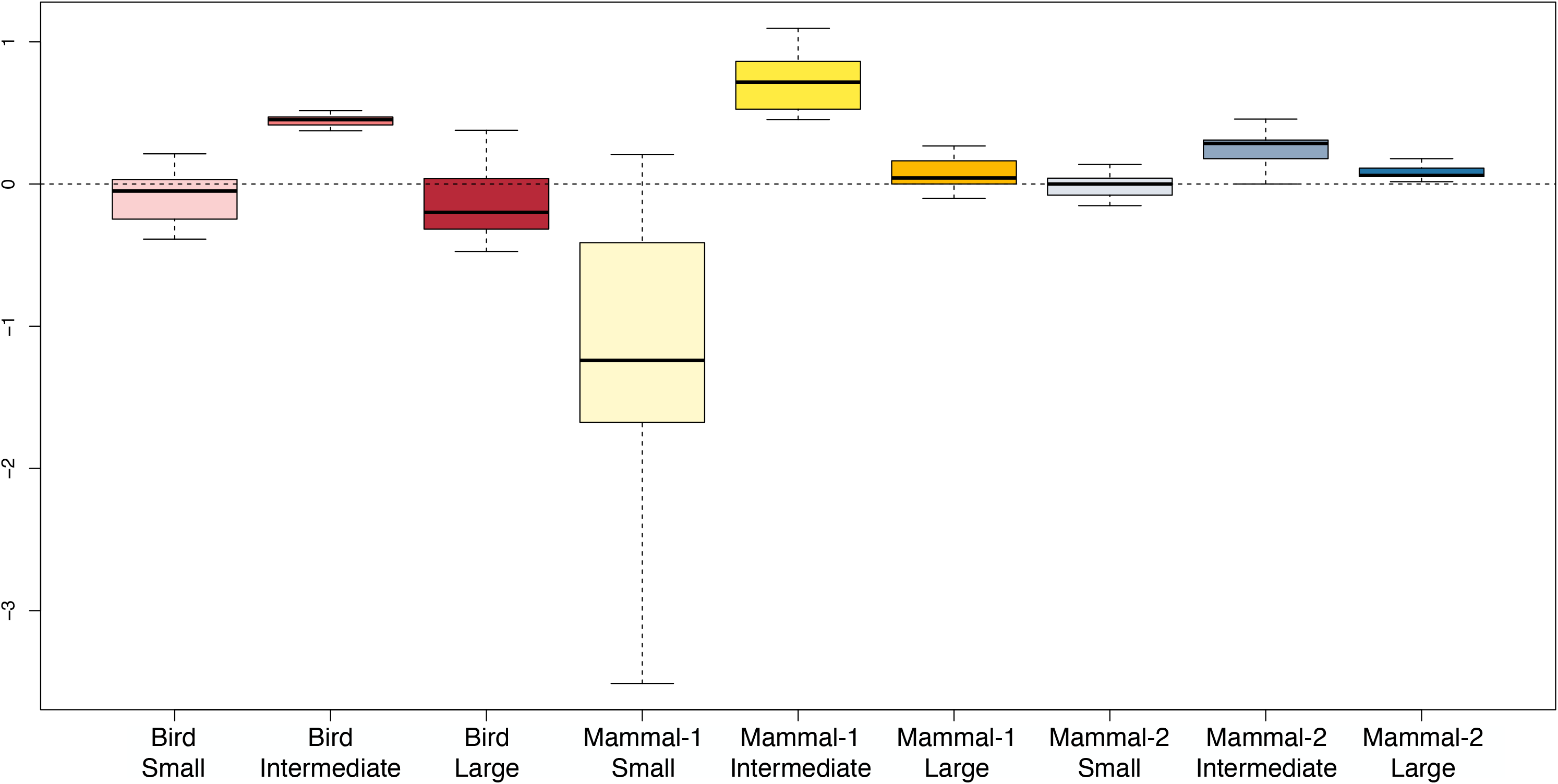
Net diversification of each combined fruit colour and range size group. Boxes show the distribution of net diversification rate values from the 30 Multi-State Speciation and Extinction models created for each of the 30 phylogenetic trees. Outliers are excluded for visibility. Each colour group is assumed to be primarily dispersed by some animal group: the first colour group by birds, the second and third by mammals.

## 4 Discussion

Fruit colour is tightly linked to dispersal and therefore to geographic range in plants. Fruit colour determines what frugivores consume which fruits and thereby disperse seeds, which together influences plant evolutionary rate (Onstein et al., 2017; Sales et al., 2021). Here we combined macroecological and macroevolutionary approaches to determine the impact of fruit colour (and hence frugivore seed-dispersal interactions) on palm range size and diversification rate. We tested two hypotheses: That (1) fruit colours attracting frugivorous bird dispersers have larger range size due to high dispersal ability. That (2) the distribution of diversification rate at different range sizes differ between fruit colour groups associated with different disperser groups, and that diversification rate would be higher for palms with intermediate range size.

Our findings show that both of the mammal-dispersed fruit colour groups have greater proportions of large-ranged species than the bird-dispersed ones. Therefore we reject our first hypothesis. There was a difference in whether small or large ranges better facilitate speciation between the bird-dispersed fruit colour group and both the mammal-dispersed fruit colour groups. For the bird-dispersed group, small ranges were associated with higher net diversification than large ranges, while for both mammal-dispersed groups, large ranges were associated with higher net diversification than small ranges. We also found that the highest diversification rate was associated with yellow fruits (i.e. mammal-1-dispersed fruit colour group), followed by black, purple, red, orange, and white fruits (i.e. bird-dispersed fruit colour group) and lastly brown, blue, and green fruits (i.e. mammal-2-dispersed fruit colour group). Therefore, dispersal ability alters at which range size diversification is higher, and intermediate range size was associated with high net diversification rate for all three colour fruit groups (i.e. bird, mammal-1, and mammal-2), confirming our second hypothesis.

### 4.1 Frugivore disperser ability and species range size

While a positive correlation between dispersal ability and range size is not ubiquitous (Gaston, 1998), several studies have demonstrated such an effect (e.g. Estrada et al., 2015; Faurby & Antonelli, 2018; Penner & Rödel, 2019; Sinnott-Armstrong et al., 2018). Our study did not identify such an effect in palms (Figure 2). This is assuming, as we have done, that the substantiated claim that birds generally have higher dispersal ability than mammals is true (Santos, Tellería, & Virgós, 1999; Lu et al., 2019). Our finding is that in both mammal-dispersed fruit colour groups (mammal-1 and mammal-2), large-ranged species make up the plurality of species (42.4% and 39.9% respectively; Figure 2). For bird-dispersed fruit colours, species with large range size is the smallest group (29.7%; Figure 2). It is however notable that our range size groups were defined to have an equal number of species in each, potentially constraining results.

Dispersal ability is just one of many factors that influence range size. After speciation there are multiple ways that the parent species range is divided among the two daughter species (Gaston 1998). For one, it may be asymmetrically divided to various extents. The division is determined by the process by which speciation occurred: allopatric, peripatric, parapatric, or sympatric. The speciation of a small-ranged species will invariably result in two small-ranged daughter species. Given our categorization of range size, a large-ranged species could speciate into daughter species with any initial combination of range sizes, given that a parent species could have a range large enough to fit two large ranges. Even within modes of speciation there may be different outcomes of range division depending on specific mechanisms, for example whether allopatric speciation occurs through vicariance or peripheral isolation (Gaston 1998).

In our study, birds have more small and medium-ranged species than large-ranged species. A possible reason the bird-dispersed group has a lower proportion of large-ranged species is the reduction in range size post-speciation for daughter lineages (Pigot, Phillimore, Owens, & Orme, 2010) followed by limits to range expansion post-speciation (Pigot, Owens, & Orme, 2010). A daughter species that has speciated through adapting to novel habitat may be limited to this specific habitat. If this novel habitat is geographically small, dispersal ability does not matter – the daughter lineage range size would remain small. If speciating through isolation by distance over a large range, a daughter lineage may not be limited by habitat in such a way, but could potentially expand its range to match the range of the ancestral species. High dispersal ability could hinder isolation by distance by maintaining gene flow. This means that speciation modes that occur most frequently for species with better dispersal ability, for example by ecological speciation after range expansion into novel habitat, can hinder subsequent range expansion.

If better dispersal ability does tend to correlate positively with range size, our results indicate that daughter lineages dispersed by long-distance dispersers (some birds) are limited from achieving large range by, for example, ecological factors. It is possible that there are other dispersal-range size dynamics that impact diversification, for example, high dispersal could result in very fragmented small populations that simply go extinct. Taken together, our results demonstrate the importance of other determinants of range size than dispersal ability in palms.

### 4.2 Frugivore dispersers alter range size-diversification rate relationship

Our study demonstrates that diversification varies depending on disperser group, which is mediated by fruit colour. For all groups, diversification was highest for species with intermediate size, however, palms with fruit colours associated with mammal-dispersal had higher net diversification at larger ranges than smaller ranges, while the opposite was true for fruit colours associated with bird-dispersal. Previous studies have also found an association between different fruit colours and diversification (Lu et al., 2019; Onstein et al., 2020), and that ecological interactions may alter patterns of diversification (Burin, Guimarães, & Quental, 2021). Range or migratory patterns of frugivorous birds and mammals affect at which range size diversification in palms occurs most frequently. This finding further corroborates the previously demonstrated importance of frugivore selection in shaping fruit traits (Valenta & Nevo, 2020).

What reasons might be behind our finding that bird-dispersed palms diversify more frequently at smaller ranges than at larger ranges? It could be due to greater maintenance of gene flow between bird-dispersed populations of large-ranged species. Due to the ability to traverse barriers and long-distance migratory patters, birds are more likely to return to areas colonised by migratory birds of previous years, such as islands, and thereby maintain gene flow (Viana, Gangoso, Bouten, & Figuerola, 2016).

Quaternary megafaunal extinctions may be influencing our finding of relative high number of large-ranged mammal-dispersed palms as well as the diversification at larger ranges compared to smaller. This extinction appears to have lead to a breakdown of fruit trait mammal interactions (Muñoz, Trøjelsgaard, & Kissling, 2019). Mobile large mammals that are now extinct might have facilitated large ranges. These palms would have either gone extinct (Onstein et al., 2018) or adapted to to new dispersers, largely scatter-hoarding rodents (Muñoz et al., 2019). Species previously dispersed by these extinct mammals may have reduced range sizes as well as elevated extinction rate, possibly explaining why we find that mammals have higher net diversification at larger ranges than small ranges.

### 4.3 Intermediate range size leads to higher diversification rate

Our findings support the hypothesis that intermediate range size is associated with the highest net diversification rate (Claramunt et al., 2012; López-Villalta, 2014; Figure 3). This result indicates that in palms the variables that influence geographic speciation do not favour diversification at either range size extremity. It could be multiple interacting variables that result in higher diversification at intermediate range size. An example of such a potential interaction is the tendency for speciation through isolation by distance at larger range size opposed to the diversification-hampering maintenance of gene flow due to higher dispersal ability of a large-range species.

Intermediate-ranged palms in the “mammal-1” group had highest net diversification overall, followed by the intermediate-ranged palms in the “Bird” group, and finally intermediate-ranged palms in the “mammal-2” group. The difference observed in net diversification rate between the two mammal-dispersed groups indicates some underlying factor(s) that differs between how the two mammal-dispersed groups of palms are dispersed.

The mammal-2 group is more strongly associated with mammal dispersal than the yellow fruits that comprise the mammal-1 group (Onstein et al., 2019; Knight & Siegfried, 1983; Willson & O’Dowd, 1989), the latter which is likely to have more bird-dispersal mixed with mammal-dispersal. Assuming then that yellow fruits have mostly mammalian as well as bird dispersers, why would that result in higher diversification rate? We suggest that with fewer birds dispersing these seeds, long-distance dispersal would be less frequent. The occasional bird-dispersal would allow for the establishment of isolated populations more likely to speciate (Parent, Caccone, & Petren, 2008). But as long-distance dispersal by birds is still scarce, the overall species range remains intermediate (with higher diversification rate, as we have shown). This logic may also explain why primarily bird-dispersed palms experience higher diversification at slightly smaller range sizes than the other two categories. If bird-dispersed, a smaller range combined with high dispersal ability results in a greater potential for colonising new areas and thereby diversifying (Lester et al., 2007). Range size that leads to higher diversification is larger for mammal-1 fruits than for the other two groups (Figure 3). This may indicate that gene flow is lower over the extremes of each of these large ranges than in the other two groups (and assuming no sympatric speciation or small-scale parapatry).

There may also be environmental factors that influence palms with yellow fruits specifically. An effect of fruits being located in understory habitat leading to higher speciation in palms has been suggested (Onstein et al., 2017). Yellow-fruited plants do not appear more frequently in understory habitats according to PalmTraits, but some other environmental factor that may be typical of yellow fruits may be influencing speciation. Lu et al. (2019) speculated that high diversification rate may be a result of the presence of fruits in habitats associated with high diversification. Previous studies have also indicated an effect of fruit size on speciation rate (Onstein et al., 2017; Lim et al., 2020). Some other fruit trait more common in yellow fruit than other colours may influence speciation rate and dispersal.

Our results stand in contrast with studies that have identified high diversification for species with large range size (e.g. Gohli & Jordal, 2017). Our results may contrast with studies that use animals as a study system due to the difference in dispersal mechanism by animals and plants that spread through endozoochory. Animals disperse by the movements of individuals within the same species, which should therefore be relatively similar. Plant-frugivore interactions are highly complex (Bascompte & Jordano, 2007). The offspring of a single tree could be dispersed by many different frugivore species with varying mobility, foraging, and movement patterns, even within the major groups in our study (birds, mammals). Varying spatial effects on diversification between animal dispersal and plant-frugivore dispersal should therefore be expected. Another effect resulting from a fundamental difference between plant and animal dispersal, such as the interplay of pollination and seed dispersal in plants (Epperson, 2007), could also be leading to these differing results.

## 5. Conclusion

Our study demonstrates that dispersal ability does not correlate positively with range size in palms. We find striking variation in diversification rate between range size categories for species with different fruit colors, indicating an effect on diversification by frugivore trait selection. The relationships between range size and diversification for different disperser groups are not clearly delineated. This highlights the importance of weighing the combination of factors that influence range size for the particular species of interest rather than assuming a general rule of diversification-range size relationship is applicable. Finally, our results show that diversification rate is highest for intermediate range sizes. This indicates that opposing factors, such as isolation by distance and gene flow, are relatively balanced in palms, or at least do not favour the smallest and largest range sizes. Dispersal is tied to geographic speciation, and therefore plays an important role in diversification. Given the role of fruits for plant dispersal it is important to examine fruit traits that influence dispersal and thereby diversification.

## ACKNOWLEDGMENTS

This work was supported by the Biodiversity and Ecosystem Services in a Changing Climate Strategic Research Area at the University of Gothenburg. C.C.S. was supported by Coordenação de Aperfeiçoamento de Pessoal de Nível Superior, Brazil. S.F. was supported by the Swedish Research Council (2017-03862); C.D.B. and M.F.T.J. were supported by the Swedish Research Council (2017-04980).

## AUTHOR CONTRIBUTION

C.D.B. conceived the ideas and all authors together designed the methodology; A.H., M.F.T.J., and S.F. collected the data, A.H. analysed the data and led the writing of of the manuscript. All authors contributed critically to the drafts and gave final approval for publication.

## DATA AVAILABILITY

The palm phylogeny used in this paper is from Faurby et al., (2016) (https://doi.org/10.1016/j.ympev.2016.03.002) with some changes. The following changes have been made to the updated version 1.1 from 2018 used in our study: The Iriarteeae was reananalysed, all at one time based on data from Bacon et al. (2016). It was rooted internally with *Iriartea*, and *Dictyocaryum* was set as sister to remaining genera based on the same source. The clade comprising *Balaka*, *Veitchia*, *Adonidia*, *Jailoloa*, *Manjekia*, and *Wallaceodoxa* was reanalysed based on Heatubun, Zona, & Baker (2014) and on Alapetite, Baker, & Nadot (2014). It was rooted with *Adonidia* as sister to the rest. *Sabinaria* was added as sister to *Itaya* based on Bogotá-A et al. (2015). The taxonomy was then updated.

The palm fruit colour and geographic occurrence data used for the analyses is publicly available in Kissling et al. 2019 (https://doi.org/10.6084/m9.figshare.9766919) and Global Biodiversity Information Facility (GBIF; ttps://doi.org/10.15468/dl.rjmqfy), respectively. The code for cleaning the data is available on GitHub (https://github.com/mftorres/palm_leaf/blob/master/notebooks/GBIF_variables_data_prep.ipynb)

## BIOSKETCH

Adrian Hill is a PhD student at the University of Gothenburg. His research interest is to better understand the evolution of plant diversity through genomic and macroevolutionary studies.

The Gothenburg Global Biodiversity Centre is hosted at the University of Gothenburg and provides a platform to better understand and communicate biodiversity sciences among its 17 partner organisations around Western Sweden. The centre’s goal is to better understand biodiversity and address issues related to the ongoing biodiversity crisis to build a biodiversity-positive future.

