## Appendix S1 for "Fruit colour and range size interact to influence diversification"

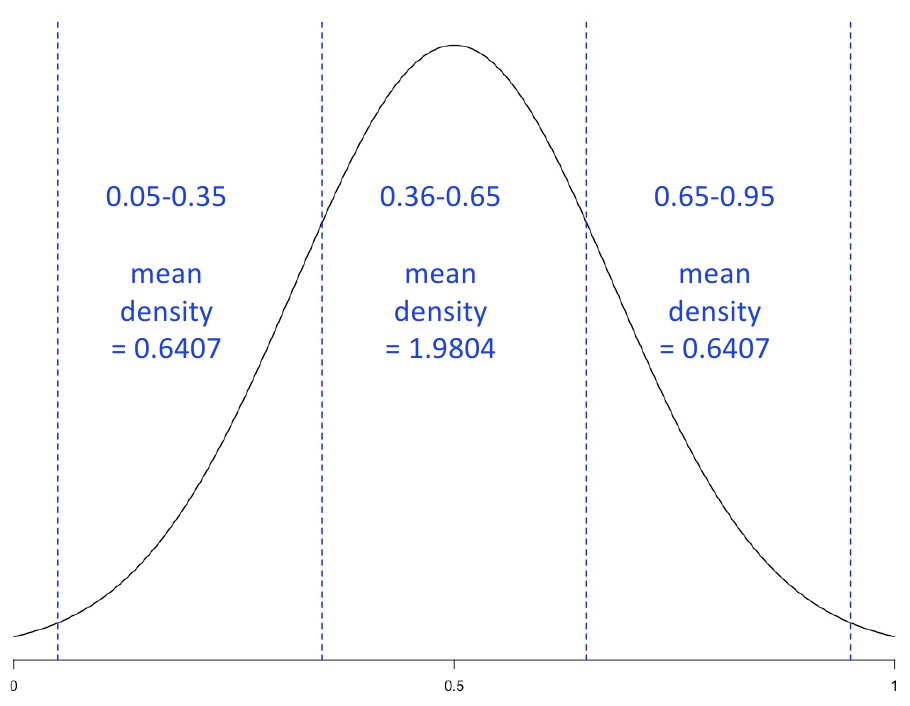
Figure S1.1. Example graph to illustrate our suggested explanation regarding the distribution of net diversification values for range size categories within each fruit colour group. We suggested that net diversification roughly follows a normal distribution based on the distribution of net diversification values within colour categories. Net diversification rates are here represented by the bell curve. Three categories are split by the blue lines, which in our study are the small, intermediate, and large range size categories. Mean density values represent net diversification. In this scenario, small and large-ranged species have the same net diversification rates, meaning that net diversification is maximised at exact intermediate.

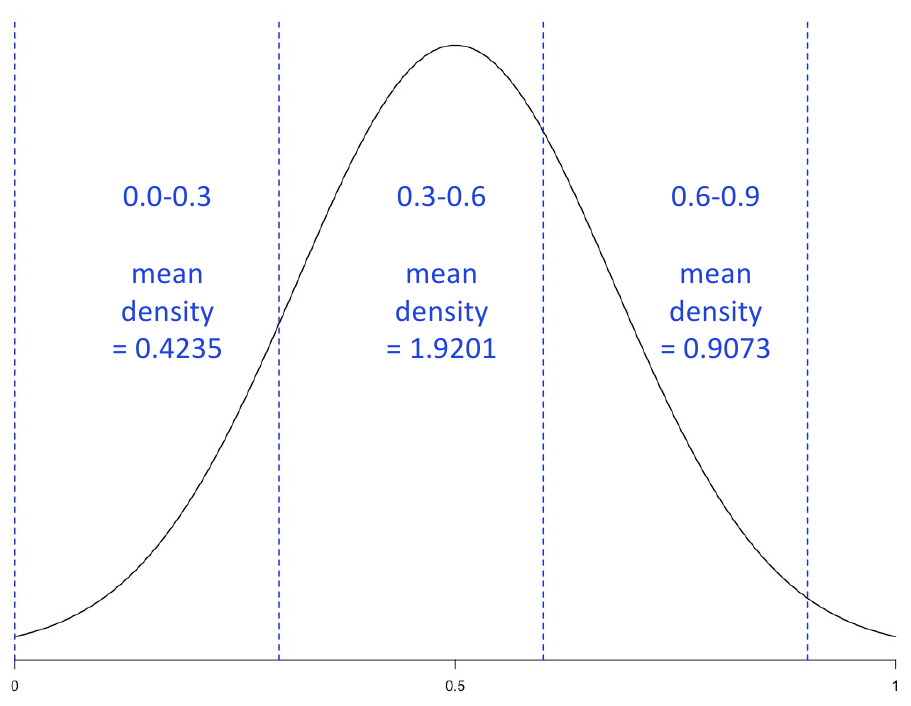
Figure S1.2. Example graph to illustrate our suggested explanation regarding the distribution of net diversification values for range size categories within each fruit colour group. We suggested that net diversification roughly follows a normal distribution based on the distribution of net diversification values within colour categories. Net diversification rates are here represented by the bell curve. Three categories are split by the blue lines, which in our study are the small, intermediate, and large range size categories. Mean density values represent net diversification. In this scenario, large-ranged species have higher net diversification than small-ranged species. This means that the range size at which net diversification is maximised is slightly higher than intermediate.

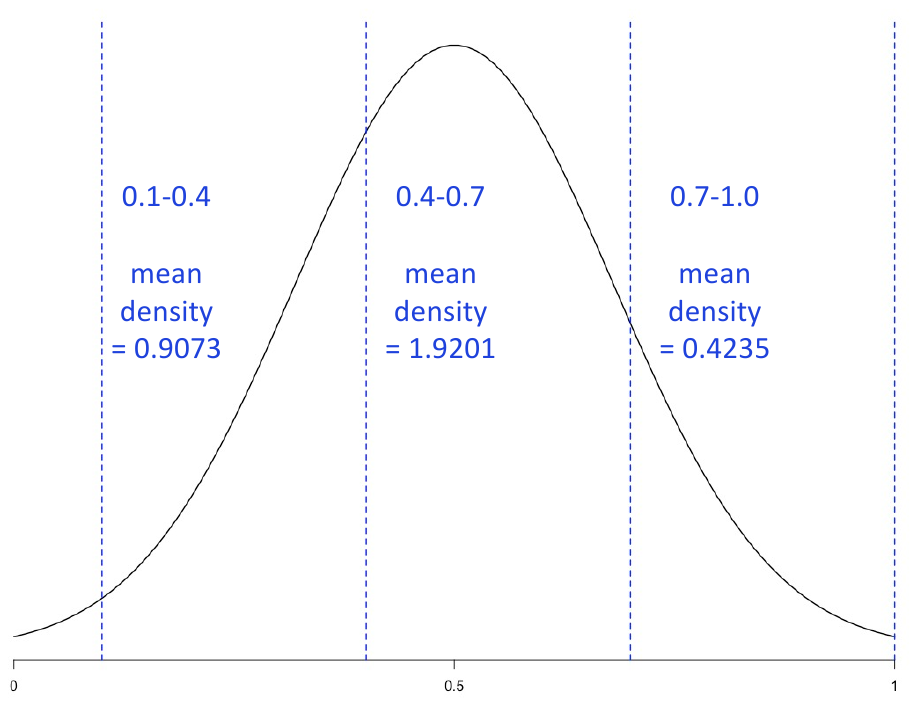
Figure S1.3. Example graph to illustrate our suggested explanation regarding the distribution of net diversification values for range size categories within each fruit colour group. We suggested that net diversification roughly follows a normal distribution based on the distribution of net diversification values within colour categories. Net diversification rates are here represented by the bell curve. Three categories are split by the blue lines, which in our study are the small, intermediate, and large range size categories. Mean density values represent net diversification. In this scenario, small-ranged species have higher net diversification than large-ranged species. This means that the range size at which net diversification is maximised is slightly lower than intermediate.

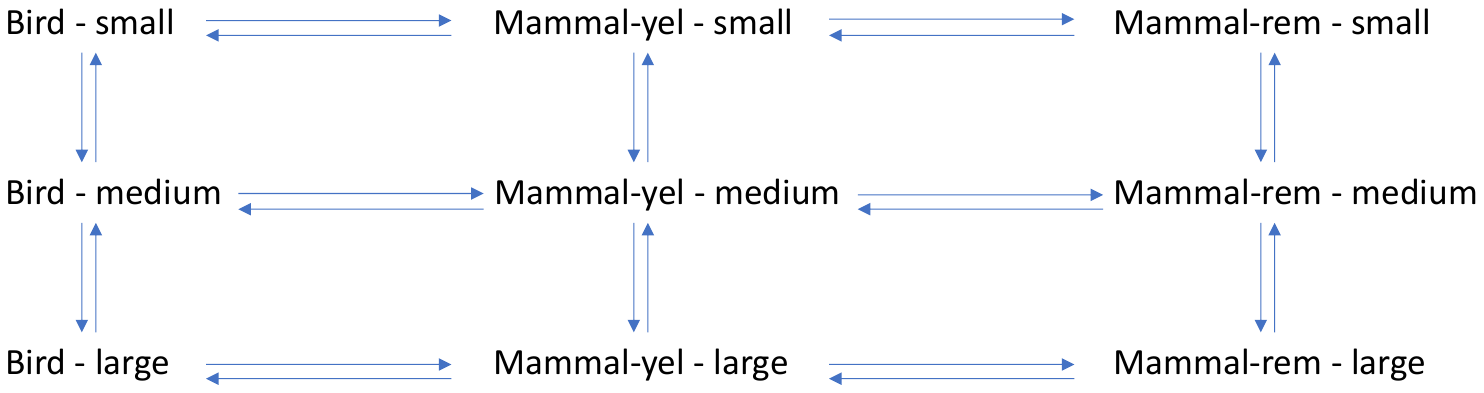
Figure S1.4. Transitions that were allowed in the nine-category Multi-State Speciation and Extinction models.
